# The tissue specificity of cancer genes

**DOI:** 10.1101/2025.09.21.677175

**Authors:** Jiayi Chen, Ryan L Collins, Kevin M Haigis

## Abstract

Defining genes that are somatically mutated in different cancers is a central goal of cancer genetics. Nevertheless, traditional definitions of “driver” genes based on pooled mutation frequency are biased toward common cancer types and tend to overlook genes that might be specific to rarer subtypes. As such, our understanding of the compendium of cancer genes is incomplete. In this study, we developed a statistical framework that defines genes enriched for functional somatic mutations in primary cancers to quantify incidence and tissue specificity for each gene. By applying this framework to the AACR GENIE v18.0 dataset, which comprised over 1.15 million mutations identified in 146,394 patients spanning 265 histologic subtypes, we identified a total of 95 genes significantly mutated in at least one subtype. We mined this dataset to derive tissue specificity scores for all 95 genes, demonstrating that tissue specificity in cancer is the norm, not the exception, for nearly all genes. We interrogated these new tissue specificity scores to reveal that oncogenes with restricted expression across normal tissues tend to exhibit higher tissue-specific mutation patterns in cancer. In contrast, tumor suppressors were often ubiquitously expressed irrespective of their underlying tissue specificity, which was partially correlated with differentially expressed compensatory genes and pathways in mutationally permissive tissues.

**Significance Statement:** We present a statistically robust, subtype-aware framework to identify tissue-specific cancer driver genes and uncover their underlying biology. Our findings reveal that tissue-selective oncogenicity arises from expression constraints and tissue-restricted compensatory pathways. These findings provide a systematic, well-powered resource on mutational significance, tissue tropism, and biological consequences for the cancer research community.

## Introduction

Identifying cancer driver genes, defined as genes wherein somatic alterations confer a selective growth advantage to tumor cells, is fundamental to understanding tumor biology and guiding therapeutic development. Large-scale efforts such as The Cancer Genome Atlas (TCGA) have profiled thousands of tumors with a variety of genomic technologies and many sophisticated methods have been proposed for nominating mutational driver genes (1, 2). As a general rule, most approaches typically define driver genes based on mutational recurrence across a pooled set of tumor samples grouped by histology or tissue-of-origin. However, this pooled approach introduces a key limitation: genes that are commonly mutated in the common cancer types that are well represented in these studies, such as lung or breast cancer, tend to dominate the resulting lists of cancer driver genes. This bias is driven by relatively modest sample sizes for less common cancers in most pan-cancer cohorts, leading to low statistical power and a high false negative rate for driver gene detection in rarer cancers. As a result, genes that play important roles in less common cancer subtypes may be underappreciated, limiting our understanding of context-specific oncogenic mechanisms, as well as the extent of tissue selectivity for each mutational driver gene across all human cancer types, irrespective of disease prevalence.

Regardless of this limitation, numerous studies have proposed that tissue specificity is in fact quite common across cancer genetics. While a few genes such as *TP53* are broadly mutated, most oncogenes and tumor suppressors show restricted distributions of somatic mutations across tissues, shaped by underlying transcriptional, epigenetic, and proteomic states (3). For example, the flagship TCGA pan-cancer driver catalog reported that 142 of 258 driver genes (55%) were associated with just one cancer type, whereas only a handful of genes, such as *TP53, PIK3CA, KRAS, PTEN*, and *ARID1A*, exhibited broad distributions across a majority of cancer types (2).

Building on this perspective, we aimed to better understand the extent of tissue specificity among functional somatic mutations across as broad a spectrum of human cancers as possible. To accomplish this, we developed a subtype-aware statistical framework that evaluates somatic mutation enrichment within each histologic subtype and applied this framework to the largest publicly available pan-cancer clinicogenomic dataset, AACR GENIE v18.0 (4). After rigorous curation and quality control, our final analysis included somatic mutation data for 590 genes in 146,394 primary tumor samples spanning 265 histologic subtypes. We identified 95 genes with study-wide significant enrichment for functional mutations in at least one cancer subtype and subsequently defined a new metric to quantify how selectively each gene is mutated across cancer types while accounting for subtype-specific differential power, mutation frequencies, and other technical confounders. This approach enabled a more balanced detection of cancer drivers across the full spectrum of human malignancies, which represents a foundation for studying tissue-restricted oncogenic processes, and especially for processes relevant to less common cancers.

## Results

### Statistically robust definitions of somatic mutation frequencies across histologic subtypes

The first goal of our study was to quantify subtype-specific somatic mutation frequencies across a broad range of genes and histologic subtypes. To maximize our power for subtype-aware analyses, we curated somatic mutation data from AACR Project GENIE (v18.0). We excluded all metastatic samples due to their distinct mutation profiles at secondary sites. We also excluded subtypes with extremely limited power even at the large overall sample size of GENIE; specifically, we did not consider subtypes represented by fewer than 0.1% of the cohort (≤151 samples). Lastly, we identified and excluded likely misannotated samples (Fig. S1). After filtering, the final cohort included 146,394 non-metastatic tumor samples—of which 651 (0.44%) were benign—spanning 265 distinct histologic subtypes and 31 tissue types according to the established OncoTree ontology (5) (Fig. 1A; Dataset S01).

**Figure 1.**
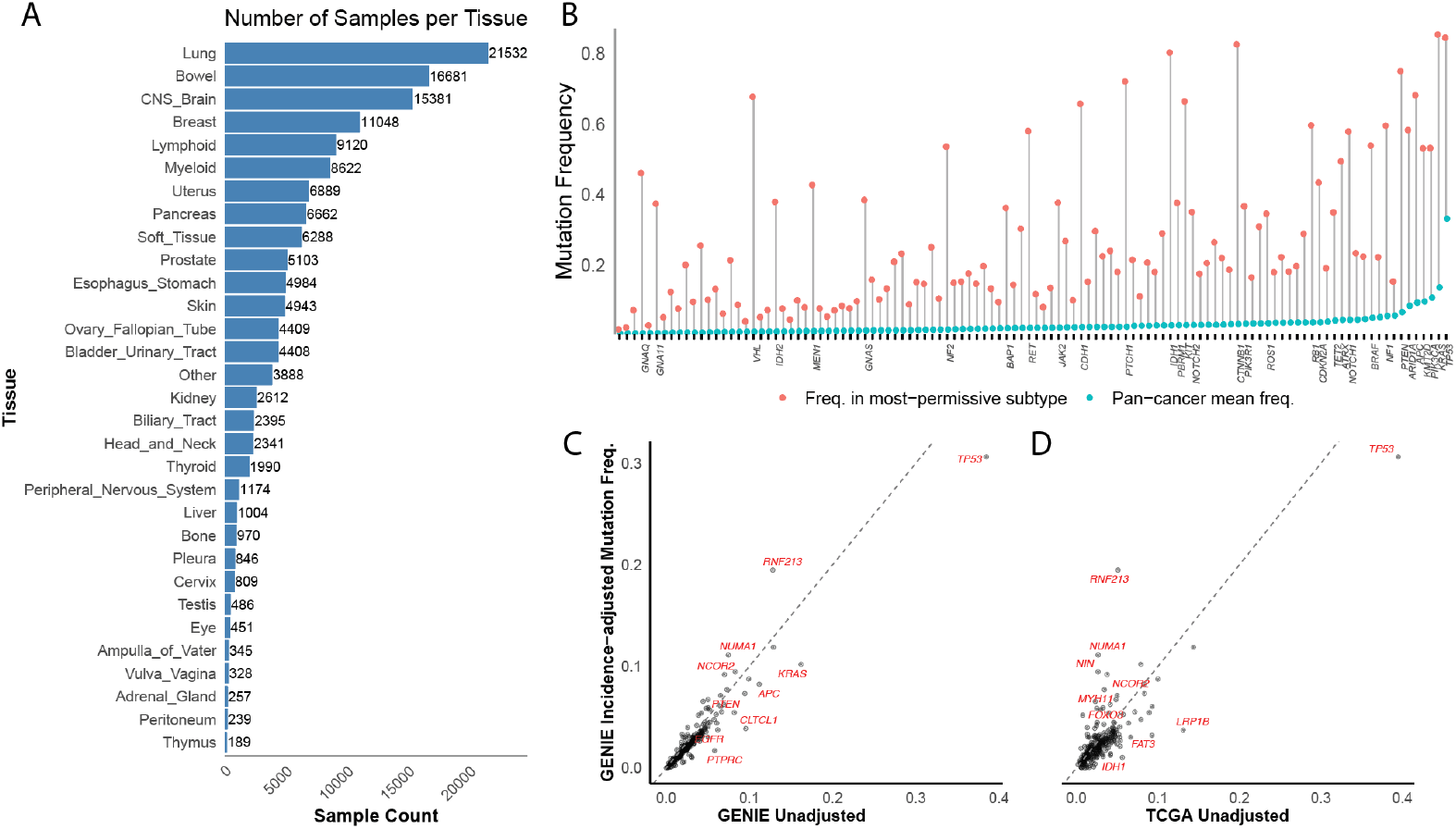
Mutational properties of the AACR GENIE v18.0 dataset after our *post hoc* curation and quality control steps. (A) Count of unique primary tumor samples included in our final analysis, stratified by major tissue (top-level OncoTree layer). (B) Many genes exhibited large discrepancies between their overall pan-cancer mutation frequency versus their highest subtype-restricted mutation frequency. Shown here are the 112 IntOGen-annotated driver genes with adequate sequencing coverage in GENIE (≥80% of patients sequenced, sequenced ≥1000 times in at least two centers, and present in ≥10% of cancer types). For each gene, a gray line segment connects its average pan-cancer mutation frequency to the mutation frequency observed in its most permissive tissue subtype. Genes are ordered along the x-axis by increasing pan-cancer rate. Genes exhibiting an exceptionally large discrepancy between their pan-cancer and subtype-restricted mutation frequencies (absolute difference >0.30) are annotated with labels on the axis. (C-D) SEER-adjusted GENIE mutation frequencies (y-axis) plotted against pan-cancer unadjusted mutation frequencies from (C) GENIE and (D) TCGA. See Methods for adjustment procedure.

Essentially all tumor samples in the AACR Project GENIE dataset were profiled using targeted somatic DNA sequencing assays (*e*.*g*., gene panel tests). Across all 146,394 samples, a total of 19,324 genes were targeted for sequencing in at least one sample; however, very few genes (364/19,324; 1.88%) were sequenced in a majority (>50%) of the overall GENIE cohort, leading to widespread differential missingness per gene across patient recruitment centers, sequencing assays, and cancer types. To account for this considerable technical heterogeneity, we curated a subset of 590 genes that had mutation frequencies high enough to rule out near-zero frequencies with 90% confidence in at least one cancer subtype (Dataset S02). To capture functional heterogeneity in somatic alterations, we assessed mutation enrichment separately for loss-of-function (LoF), missense, and combined LoF or missense consequences.

We first wanted to establish an initial assessment of discrepancies between pan-cancer and tissue-specific mutational frequencies in this dataset. To achieve this, we focused on a set of 112 genes reported as cancer drivers in the IntOGen database that also met strict coverage criteria in GENIE (≥80% of patients sequenced, ≥1000 times in at least two centers, and present in ≥10% of cancer types) (6). For each gene, mutation frequencies were calculated using the combined frequency of LoF and missense mutations for two viewpoints: (1) the mean mutation frequency across all histologic subtypes, and (2) the mutation frequency for each gene’s most permissive histologic subtype, defined as the subtype with the highest lower bound of the 90% confidence interval (CI) for that gene’s mutation frequency, which was necessary to account for variable sample sizes between subtypes (Fig. 1B). These analyses revealed the average mutation frequency for most genes (109/112; 97.3%) was below 10% in the pan-cancer setting across the pooled cohort, with 15.2% (17/112) being mutated in fewer than 1% of all samples, underscoring the sparsity of mutations in most genes when aggregated naively across all cancer types. In contrast, all genes reached mutation frequencies greater than 5% within at least one specific subtype. For example, *GNAQ* had an overall pooled mutation frequency of just 0.59% across all samples in GENIE but reached a mutation frequency of 46.1% (90% CI = 40.5–51.8%) in uveal melanoma, reflecting strong tissue specificity. Similarly, *VHL* was mutated in 1.06% of all samples but reached 67.8% (90% CI = 65.4–70.0%) in clear cell renal cell carcinoma, where it is a well-established subtype-specific canonical driver gene, while its mutation frequency remained low in non-clear cell renal cancers (5.7%; 90% CI = 4.5–7.2%). These results emphasize the importance of applying a subtype-aware viewpoint in analyses of somatic mutational enrichments, which can unmask potential subtype-restricted drivers that might be obscured by pooled pan-cancer or gross organ-level summaries.

### Estimating population-level gene mutation frequencies by combining molecular profiling and epidemiologic data

The overall frequency of functional somatic mutations in each gene across all human cancers is a critical statistic for prioritizing diagnostic, preclinical research, and therapeutic development efforts. In practice, such global mutation frequencies have been reported directly from large-scale cancer genomic datasets like TCGA and GENIE, where certain rare cancer types are overrepresented relative to their true population prevalence (*e*.*g*., sarcomas, adrenal tumors) while common, typically older-onset cancers are sometimes underrepresented (*e*.*g*., prostate). These discrepancies largely reflect clinical sequencing practices, as patients with more aggressive, treatment-resistant, or recurrent tumors are more likely to undergo clinical genomic profiling.

We aimed to correct for these biases and more accurately approximate the real-world impact of gene mutations across cancer types. To accomplish this, we projected the observed mutation frequencies per gene in GENIE through population-level cancer incidence estimates from NCI SEER (Dataset S03) (7). Overall, pan-cancer mutation frequencies per gene were broadly concordant between SEER-adjusted and unadjusted models (R^2^=0.7929; P<10^-20^; Pearson correlation test), but notable discrepancies were observed for certain genes enriched in subtypes with disproportionate representation in sequencing datasets (Fig. 1C). For example, *KRAS* exhibited an unadjusted mutation frequency of 16.2% across all primary tumor samples in GENIE v18.0 but dropped to 10.2% after SEER adjustment, reflecting its relative enrichment among aggressive cancers that are more likely to undergo genomic profiling as a part of their clinical management. Similarly, *APC* decreased from 11.2% to 8.2%, and *TP53* decreased from 38.5% to 30.6%. These shifts illustrate how raw mutation frequencies derived from pan-cancer sequencing projects may overstate the abundance of mutations prevalent in frequently sequenced tumor types, while SEER-adjusted frequencies offer a more representative approximation of the true population-level mutational burdens across all unselected human cancers.

We also noted a handful of genes that exhibited the opposite trend to genes like *VHL, KRAS*, and *APC* (Fig. S2). For example, *AHNAK2* exhibited a high pan-cancer mutation frequency in GENIE (16.05%) but substantially lower SEER-adjusted frequency (2.74%), despite being enriched in a high-incidence cancer, breast cancer. Closer inspection revealed that *AHNAK2* was sequenced almost exclusively (96.6%) in breast tumors in GENIE. Such cases, where a gene is sequenced only in select tissues, can inflate unadjusted mutation frequencies and deflate incidence-adjusted estimates. These effects highlight how gene panel design, often informed by prior knowledge or assumptions about tissue relevance, can introduce ascertainment bias.

Although our study was focused on AACR Project GENIE, we also wanted to explore these similar biases in gene mutation frequencies in TCGA, the most widely referenced pan-cancer clinicogenomic dataset (Dataset S03). As with GENIE, the mutation frequencies for most genes were well-correlated with their projected “true” population-wide frequencies (R^2^=0.3142; P=5.2×10^-20^; Pearson test). However, a handful of genes, like *RNF213* and *LRP1B*, stood out as conspicuous outliers. Both are exceptionally long genes (97th and >99th percentiles, respectively, for gene length across all protein-coding human genes), making them more susceptible to inflated mutation frequencies due to elevated passenger background mutations (Fig. S3). Thus, observed mutation frequencies in large-scale sequencing efforts may reflect both biological factors (*e*.*g*., gene length) and clinical sequencing biases, which can be overcome by careful normalization of raw sequencing data and population-level adjustment to better contextualize mutation frequency estimates across subtypes.

### Tissue specificity is nearly ubiquitous across cancer gene mutations

Having quantified the pan-cancer and subtype-specific mutation frequencies for 590 genes across 265 cancer types, we next wanted to statistically determine which gene-subtype pairs exhibited enrichments of mutations significantly above the overall background rate of functional mutations within each subtype. We designed and implemented a statistical framework to perform these analyses and assessed all results at study-wide Bonferroni-corrected significance (P <1.60×10^-7^). An important aspect of this statistical framework was a *post hoc* empirical Bayes strategy to determine a minimum fold-enrichment of somatic mutations (>10-fold; see Methods) that distinguished established, biologically meaningful driver genes from genes that were subtly enriched for somatic mutations but only achieved statistical significance due to very large sample sizes for highly abundant cancer types (*e*.*g*., lung, breast, colorectal). This *post hoc* fold-change-based filter dramatically improved comparability between rare and common subtypes by accounting for the greater power to detect significantly elevated mutational enrichments in well-represented subtypes vs. rare subtypes; for example, the interquartile range of counts of significant genes per subtype reduced from 17 to 5 after this *post hoc* adjustment (Fig. S4)

In total, we detected 487 significant gene-subtype pairs involving 95 distinct genes enriched >10-fold in at least one subtype (Fig. 2A). Reciprocally, we identified at least one significant gene for most subtypes (146/265, 55.1%; Fig. S5). Only two cancer subtypes, dedifferentiated liposarcoma and ependymoma, showed no mutational enrichments despite both having >100 patient samples and sufficient sequencing depth in ≥30% of genes (177/590) to reach a retrospective power >0.5, based on the observed number of mutation calls. This lack of enrichment is consistent with their known low mutational burden (8). To confirm that our framework recapitulated the known patterns of cancer gene function, we next examined mutation consequences by genes’ annotated roles in cancer, finding that tumor suppressors were more frequently enriched for LoF mutations compared to oncogenes (P<10^-20^), consistent with their respective inactivation- and activation-driven roles in cancer (Fig. 2B). Other than *TP53*, which was a pronounced outlier that displayed widespread enrichment across virtually all tissues, most other cancer genes were enriched for mutations in only a limited number of cancer subtypes. Furthermore, even within the same tissue, mutation frequencies for many genes varied substantially across histologic subtypes (Fig. 2C). As mentioned above, *VHL* was highly mutated in clear cell renal cell carcinoma but nearly absent in other kidney cancers. Similarly, *RB1* was enriched exclusively in small cell lung cancers (mutation frequency of 42.1%; 90% CI = 38.8–45.4%), but no other lung histologies. In contrast, some genes, like *KRAS*, exhibited comparatively broad enrichment across subtypes within a given tissue or organ but nonetheless did exhibit notable variation in mutation frequencies among subtypes. *KRAS* was mutated in >39% of samples across five of seven subtypes of bowel cancer, reaching 47.4% in mucinous adenocarcinomas of the colon, but was rarely mutated in two subtypes: anal squamous cell carcinomas (1.5%) and well-differentiated small bowel neuroendocrine tumors (~0%). These examples underscore that many genes exhibit sharp mutation specificity not only across—but also within—tissues.

**Figure 2.**
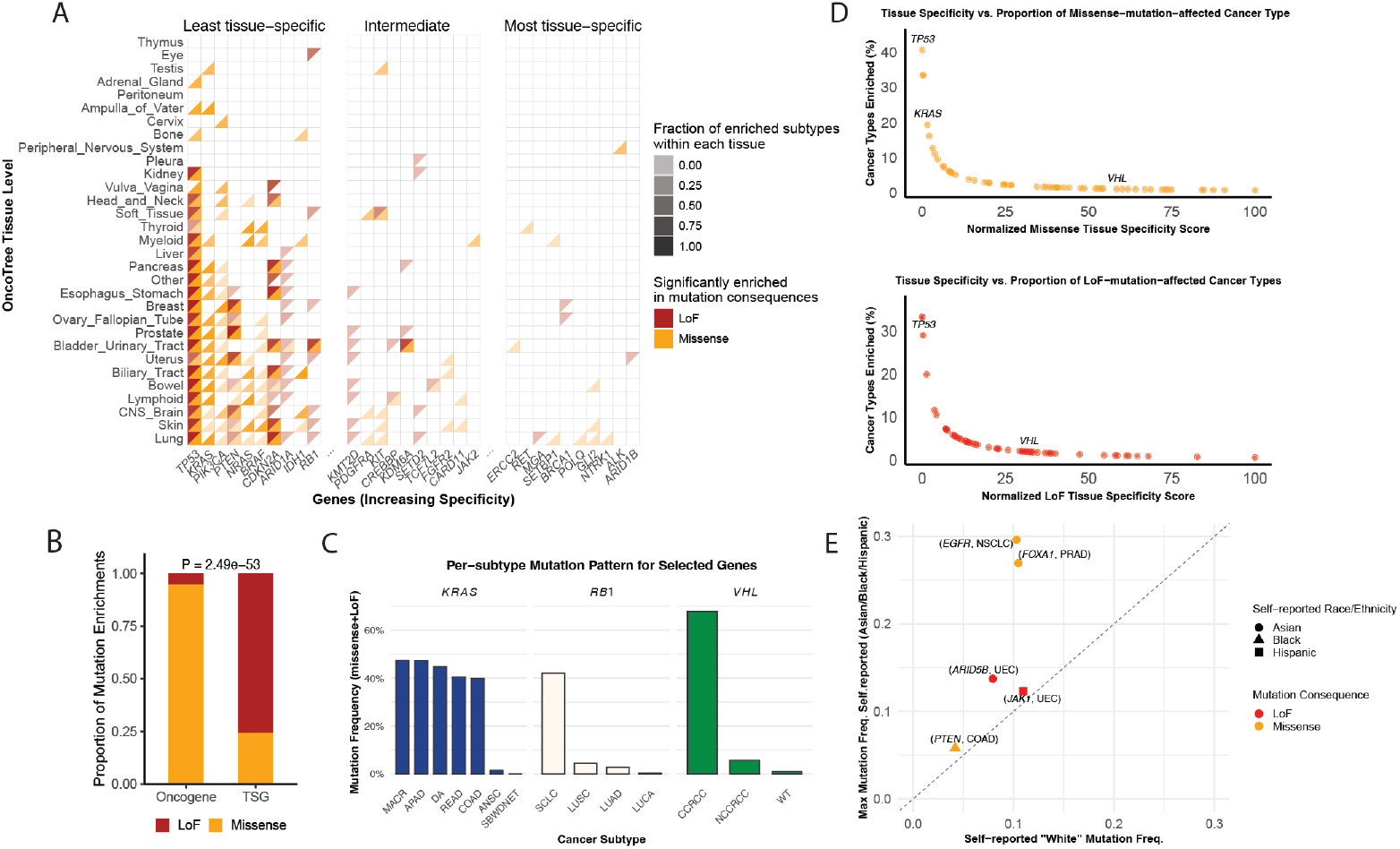
Landscape and specificity of gene mutation enrichments across cancers. (A) Mutation enrichment landscape across the 31 top-level tissues in OncoTree. Each tile represents a single gene–tissue pair; colored tiles indicate pairs with statistically significant enrichment (P<1.60×10^-7^). Lower-right orange triangles denote enrichments of missense mutations; upper-left red triangles denote enrichments of LoF mutations. Opacity reflects the proportion of mutationally enriched histologic subtypes within each tissue. Genes are ordered by increasing tissue specificity; tissues are clustered by mutational profile similarity. No fold-change filter was applied for this visualization.(B) Virtually all (94.8%) of the mutational enrichments we observed among established oncogenes corresponded to missense mutations, whereas most associations involving established tumor suppressor genes (TSGs) were composed of loss-of-function (LoF) mutations. (C) Subtype-level mutation frequencies for representative genes in permissive tissues: *KRAS* in bowel, *VHL* in kidney, and *RB1* in lung. These examples illustrate how subtype-level resolution can reveal important inter-subtype differences within overall tissue-level enrichments. (D) Relationship between our tissue specificity scores and overall pan-cancer mutation frequencies per gene. Each point represents a gene that was significantly enriched for mutations in at least one cancer. (E) We identified five genes that were significantly enriched for mutations in self-reported non-White patients as compared to White patients within the same histologic cancer subtype.

We next wanted to systematically quantify this observed tissue specificity across all mutationally enriched cancer genes. We therefore defined a “tissue specificity score” for each gene based on the fraction of sufficiently powered cancer types in which the gene was significantly enriched (Dataset S04. gene list ordered by tissue specificity score). We computed these scores separately by mutational consequence (*e*.*g*., missense, LoF) and subsequently scaled these scores from 0 to 100, with higher values indicating greater tissue specificity. Contrary to some conventional viewpoints that many “hallmark” cancer genes are mutationally relevant across many tissues and malignancies, our analysis underscored that a surprising degree of tissue selectivity of cancer gene mutations is in fact the norm, with *TP53* being a singular rare exception as perhaps the only true “pan-cancer” driver gene (Fig. 2D). Most genes, such as *VHL* and *KRAS*, exhibited significant enrichment in a small subset of cancer types, emphasizing that colloquial discussions of a gene as a “cancer gene” must be grounded in specific tissue or histologic context, even for very deeply studied genes like *KRAS*.

### Limited evidence for race/ethnicity-specific driver genes in most cancer types

We next wanted to understand the extent to which mutational enrichments within a cancer type varied by other patient-specific factors, such as race or ethnicity. We therefore stratified our mutational enrichment analyses by patient-reported race/ethnicity labels as recorded in GENIE, which included the groups “White” (N=76,509), “Spanish/Hispanic” (N=9,472), “Asian” (N=6,502), and “Black” (N=6,365). Importantly, as germline genetic data was not available for GENIE patients, we emphasize that these groups were taken directly from the GENIE metadata and do not reflect true underlying genetic ancestry; nonetheless, they provided an initial viewpoint for exploring biases of mutation frequencies between these population.

Across all 1,579 genes with at least one somatic mutation reported in GENIE, we identified just five gene-cancer pairs that were significantly enriched for somatic mutations in non-White populations, with a significantly weaker (<10-fold) and/or non-significant (P>0.05) enrichment in White patients (Table 1). These included some known ancestry-biased genes, such as *EGFR* missense mutations in non-small cell lung cancer (NSCLC) enriched in Asian patients and *FOXA1* missense mutations in prostate adenocarcinoma (PRAD) enriched in Asian and Black patients (9–11), as well as less well-characterized examples, such as *ARID5B* LoF mutations in uterine endometrioid cancer (UEC) in Asian patients (12), *JAK1* LoF in UEC in Hispanic patients, and *PTEN* missense mutations in colon adenocarcinoma (COAD) in Black patients (13) (Fig. 2E). Of these five examples, we noted that both *EGFR* in NSCLC and *PTEN* in COAD were nominally enriched in White patients but with <10-fold enrichment. We therefore applied a *post hoc* binomial test to directly compare mutation frequencies between White and non-White groups. Four of the five genes—*EGFR* in NSCLC, *FOXA1* in PRAD, *ARID5B* in UEC, and *JAK1* in UEC—passed this *post hoc* binomial test for significant enrichment in at least one non-White group after Benjamini-Hochberg correction, confirming that the mutation frequencies in these four genes was significantly elevated in non-White patients. The fifth gene, *PTEN* in COAD, showed a slightly elevated mutation frequency in Black patients (5.8%) versus White patients (4.2%), but this difference did not reach statistical significance after FDR correction.

**Table 1.**
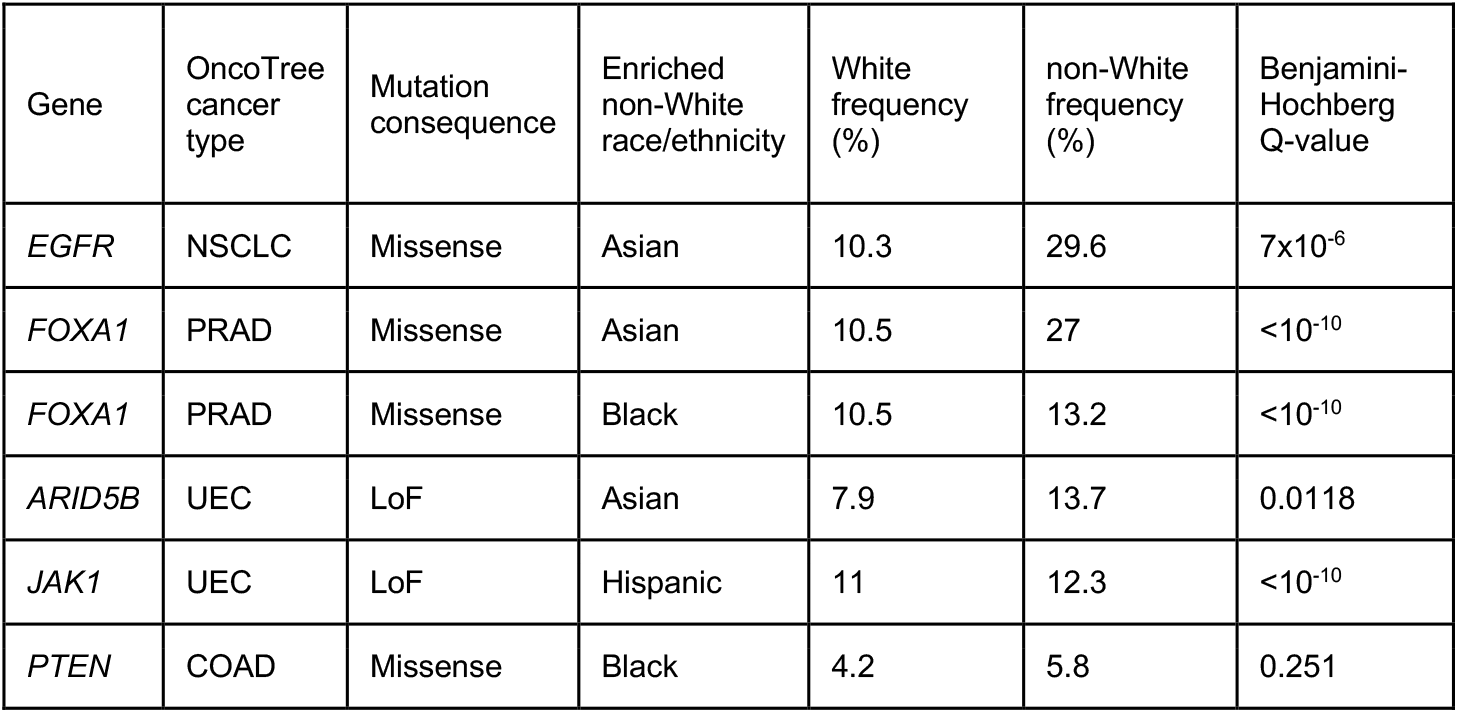
Genes with greater enrichments of mutations in self-reported non-White vs. White patients.

| Gene | OncoTree cancer type | Mutation consequence | Enriched non-White race/ethnicity | White frequency (%) | non-White frequency (%) | Benjamini-Hochberg Q-value |
| --- | --- | --- | --- | --- | --- | --- |
| *EGFR* | NSCLC | Missense | Asian | 10.3 | 29.6 | $7 \times 10^{-6}$ |
| *FOXA1* | PRAD | Missense | Asian | 10.5 | 27 | $<10^{-10}$ |
| *FOXA1* | PRAD | Missense | Black | 10.5 | 13.2 | $<10^{-10}$ |
| *ARID5B* | UEC | LoF | Asian | 7.9 | 13.7 | 0.0118 |
| *JAK1* | UEC | LoF | Hispanic | 11 | 12.3 | $<10^{-10}$ |
| *PTEN* | COAD | Missense | Black | 4.2 | 5.8 | 0.251 |

We further confirmed that each gene-cancer association was reported by at least three independent sequencing centers, and none of these genes were significantly enriched in White patients in any parent or sister/cousin cancer type in OncoTree (*e*.*g*., *EGFR* was not significantly mutated with >10-fold enrichment in any other lung cancer subtype among White patients). These checks guarded against biases that could arise if certain race/ethnicity groups were disproportionately sequenced at centers that differed in the level of detailed sample annotation they submitted to GENIE. Thus, while these analyses were constrained by comparatively limited statistical power in non-White populations, even the smallest group included thousands of patients. The fact that only five genes met our significance criteria therefore represents a small number of ancestry-specific differences. Overall, these results suggested that many driver genes discovered in predominantly White populations are likely also relevant in other racial and ethnic groups, though further studies in more diverse cohorts remain essential, especially for rare cancers where statistical power is an even greater concern.

### Gene expression in normal tissues partially explains tissue specificity of cancer mutations

Having quantified tissue specificity scores for the 95 significantly mutated genes, we next sought to investigate the biological factors that might underlie their tissue-selective mutational patterns. Specifically, we considered whether baseline RNA expression levels across normal tissues might help explain why certain genes incur somatic mutations observed in a restricted subset of cancers. We hypothesized that genes broadly expressed across many normal tissues would be permissive to somatic mutations in a wide range of cancer types, leading to lower tissue specificity scores, whereas genes with narrower expression profiles might exhibit more selective patterns of mutation enrichment across cancer subtypes.

To test this hypothesis, we obtained median gene-level RNA expression quantifications from 23 normal, non-cancerous tissues from postmortem donors from the Genotype-Tissue Expression (GTEx) Project (v10) and computed an “expression breadth” statistic for each gene, defined as the number of tissues in which expression exceeded a modest level of detectable expression (>5 transcripts per million [TPM]; Fig. S6) (14). We then assessed the relationship between expression breadth and our cancer subtype specificity scores using Spearman’s rank correlation (Fig. 3A-B). We further stratified this analysis by oncogenic mechanism, as we reasoned that proto-oncogenes and tumor suppressor genes might have different RNA expression patterns in normal tissues. For our tumor suppressor-focused analyses, we included all 54 significantly LoF-enriched genes given that most such genes (85.4%) corresponded to established tumor suppressors annotated in COSMIC (15). However, as missense mutations likely represented a mixture of damaging, LoF-like alleles and oncogenic, gain-of-function alleles, we restricted our analyses of missense mutations to known oncogenes according to COSMIC (n = 25).

**Figure 3.**
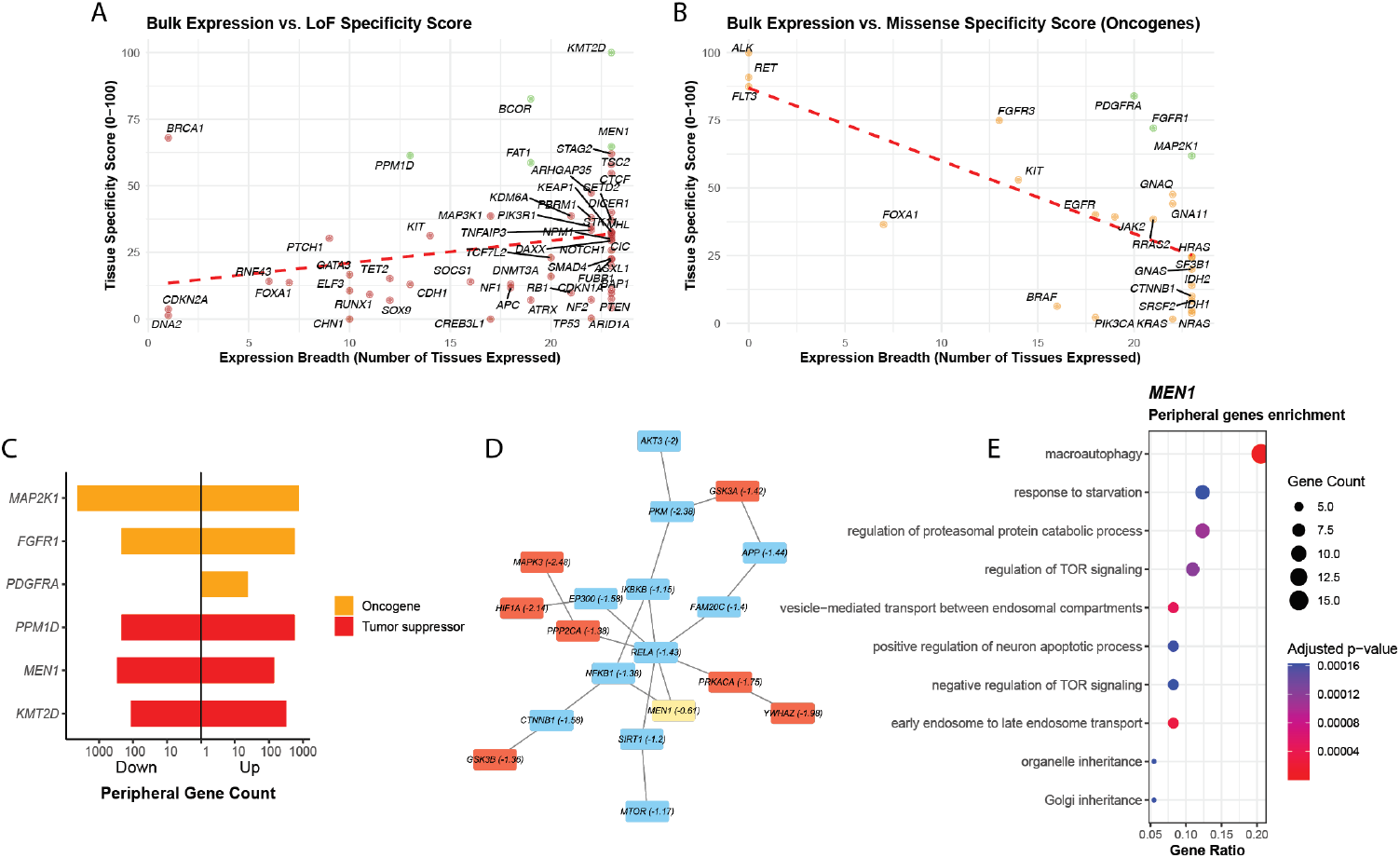
Expression breadth, tissue specificity, and pathway context of gene mutation enrichments. (A-B) Relationship between expression breadth in normal tissues and tissue specificity scores for (A) genes enriched in LoF mutations and (B) oncogenes with missense mutations. Each point represents a gene, with a dashed red line indicating the linear regression fit. Genes of interest referenced later in the text are highlighted in green. (C) Numbers of peripheral genes for three LoF mutation enriched tumor suppressors (*KMT2D, MEN1, PPM1D*; red) and three missense mutation enriched oncogenes (*PDGFRA, FGFR1, MAP2K1*; orange). Bars to the left represent downregulated peripheral genes and bars to the right represent upregulated peripheral genes. (D) PPI network centered on *MEN1*, displaying selected first-through fourth-degree branches that terminate in peripheral genes or include key pathway nodes (*MTOR, AKT*). Nodes are annotated by differential expression status: significant FDR < 5% differentially expressed genes are shown in red, and all others in blue. (E) GO enrichment analysis of *MEN1*-associated peripheral genes, showing the top 10 biological process terms with the lowest FDR-adjusted p-values. These terms converge on terms related to regulation of mTOR signaling.

Missense-enriched oncogenes exhibited a negative correlation between tissue specificity and expression breadth (Spearman ρ = -0.294, P=0.0309), consistent with the hypothesis that mutations preferentially conferred selective advantages in tissues where the proto-oncogene was endogenously expressed, allowing the oncogenic mutant allele to be expressed in precancerous cells. We noted that three genes—*ALK, RET*, and *FLT3*—were extreme examples of this trend, exhibiting high missense subtype specificity and an extremely narrow spectrum of normal tissue expression; curiously, all three of these genes are also known to be involved in recurrent oncogenic fusion rearrangements in various cancer types (16–19). To confirm this overall correlation was not driven by these three genes, we re-fitted a linear model after removing *ALK, RET*, and *FLT3*, which revealed no significant differences in this overall trend (Welch’s two-sample t-test of regression coefficients, t=-0.651, P=0.518). In contrast, LoF-enriched genes showed a significant positive correlation between expression breadth and tissue specificity (Spearman ρ=0.324, P=0.0168). This pattern likely reflects the central roles of many tumor suppressor genes in cancer prevention across myriad tissue contexts, as tumor suppressors tend to be ubiquitously expressed and involved in essential cellular processes such as genome maintenance, cell cycle regulation, and apoptosis. Collectively, these analyses provided support for our overarching hypothesis that the tissue-specificity of somatic mutations in cancer is partially dependent on endogenous gene expression levels in each cancer’s tissue-of-origin, and that this relationship markedly differs between oncogenes and tumor suppressors.

### Tissue-specific compensatory pathways contribute to extreme examples of subtype-specificity

Another common feature of some tumor suppressor genes is functional redundancy with other tumor-suppressive pathways expressed in the same tissues (20). However, in specific tissue contexts where these compensatory pathways are not utilized, LoF mutations in tumor suppressor genes can be selectively advantageous. As a result, even broadly expressed tumor suppressors may exhibit strong mutational specificity. Accordingly, we noted that a handful of genes—including both oncogenes and tumor suppressors—stood out as particularly subtype-specific mutational outliers despite being nearly ubiquitously expressed in normal tissues. We hypothesized that these outliers may stem from common properties of the biological networks surrounding these genes and therefore sought to investigate the role of compensatory pathways in shaping their selective landscape across tissues. We focused on a set of six exemplar genes that exhibited broad expression across normal tissues but had strong subtype-specific mutation enrichment. For uniformity, we selected three LoF-enriched tumor suppressors (*KMT2D, MEN1*, and *PPM1D*) and three missense-enriched oncogenes (*PDGFRA, FGFR1*, and *MAP2K1*), which we refer to as “anchor” genes in subsequent analyses.

We reasoned that the mutationally permissive tissues for anchor tumor suppressors may lack compensatory mechanisms that are otherwise active in non-permissive tissues. Similarly, we hypothesized that the tissues permissive to oncogenic mutation in anchor oncogenes may have signaling networks involving these oncogenes that are uniquely primed in certain tissues to be sensitive to the molecular consequences of the oncogenic mutation. To test these hypotheses, we first identified “peripheral” genes that were differentially expressed in permissive vs. non-permissive normal tissues for each anchor gene at a false discovery rate of 5% (Fig. 3C). We reasoned that the set of all differentially expressed peripheral genes would likely represent a mixture of the compensatory pathways that could modulate the phenotypic impact of anchor gene mutations, as well as any tissue-specific processes that were functionally irrelevant for the cancer gene in question. Thus, to further place these peripheral genes in the context of known networks involving each anchor gene, we queried the ENSEMBL IntAct Molecular Interaction Database to identify physical protein-protein interactions (PPIs) between each anchor gene and its peripheral partners (21). We constructed PPI networks up to four degrees of separation from each anchor gene and performed Fisher’s exact tests to determine whether peripheral genes were significantly enriched within each anchor’s network.

This analysis revealed significant enrichment of differentially expressed peripheral genes at higher degrees of PPI connectivity (Fig. 3D). Among tumor suppressors, *MEN1* exhibited strong peripheral gene enrichment among third-degree interactors (OR=2.03, P=0.0021) and fourth-degree interactors (OR=1.36, P=0.0245), while *KMT2D* showed enrichment for third-degree interactors (OR=1.51, P=0.0158). Among oncogenes, *MAP2K1* demonstrated peripheral gene enrichment for third- and fourth-degree interactors (third-degree OR=1.17, P=0.0334; fourth-degree OR=1.23, P=6.5×10^-5^). No significant enrichment was observed among first- or second-degree interactors for any anchor gene, likely reflecting limited power due to the small number of direct interactors at lower degrees: for example, these anchor genes had a median of only 4.5 first-degree PPIs (range = 3–16). These findings support a possible model in which tissue-specific protein networks can modulate the mutational selectivity of broadly expressed genes, irrespective of their functional class.

Lastly, to characterize the specific biology implicated in tissue permissivity for each anchor gene, we performed Gene Ontology (GO) enrichment analysis of all differentially expressed peripheral genes within four degrees in each anchor gene’s PPI network. For example, *MEN1*-associated peripheral genes within the four-degree PPI showed significant enrichment for pathways related to the regulation of mTOR signaling (Fig. 3E). These peripheral genes were downregulated in pancreas, the tissue permissive to *MEN1* mutations in our analyses; indeed, *MEN1* is well-known as a canonical driver specifically in pancreatic neuroendocrine tumors (22). Within four PPI degrees, *MEN1* was connected to seven peripheral genes annotated as negative regulators of mTOR, all of which coverage on the *RELA*–*SIRT1*–mTOR axis, consistent with prior reports in pancreatic neuroendocrine tumors (23). This pattern provides a potential explanation for *MEN1*’s mutational tissue-specificity: if inhibitory components of the mTOR pathway are already repressed in pancreas relative to other tissues, then LoF mutations in *MEN1* may further weaken this regulatory axis, thereby enhancing mTOR activity and promoting tumorigenesis. Collectively, these analyses highlighted that tissue-selective oncogenesis is shaped by both intrinsic gene expression levels activity and extrinsic cellular context, each acting as a gatekeeper for mutational impact.

## Discussion

In this study, we conducted a systematic statistical survey of nonsynonymous mutation enrichments in hundreds of putatively cancer-relevant genes across most histologic subtypes of human cancers. Our analyses demonstrated that tissue selectivity is the prevailing rule among virtually all cancer mutational driver genes, with only one exception—*TP53*—showing broad, pan-cancer enrichment (3). Our work builds upon seminal prior studies from TCGA and others, which arrived at initial coarse estimates of subtype specificity based on thousands of samples and dozens of cancer types (24). For example, the TCGA consortium estimated that most cancer genes were mutated in only 2-20% of tumors (24). In our analysis of 146,394 samples spanning 265 histologic subtypes, we similarly observed that few genes are mutated at high frequency across all tumors. However, by evaluating each subtype separately, we found that many of these genes show clear enrichment within specific tissue contexts, suggesting that the intermediate mutation frequencies in pan-cancer datasets can mask prominent subtype-specific patterns. By implementing this subtype-aware statistical framework, which adjusted for many biases endemic to large, aggregated clinicogenomic cancer cohorts, we were able to document a more uniform landscape of genes with uniformly strong enrichments of nonsynonymous mutations in both common and rare cancer subtypes, including genes that would likely be overlooked in conventional pooled analyses: 11.6% of the genes we identified as significantly enriched in at least one subtype were not catalogued as drivers in intOGen (e.g., *TNFAIP3, NPM1, DNA2, H3F3A*), 7.4% were absent from COSMIC (e.g., *SOX9, ACD, TLR4, ALB*), and 6.3% were absent from both. These findings suggest that our framework may expand the current catalog of candidate driver genes by capturing subtype-restricted signals that are missed in many pan-cancer analyses.

In contrast, we were surprised to detect only five candidate gene-cancer pairs that exhibited significant mutational enrichments in self-reported non-White vs. White patients, implying that there is apparently limited variability in strong-effect mutational drivers of cancer across major race/ethnicity groups. As germline genetic ancestry is not available for the GENIE dataset, these findings need to be confirmed in future well-powered studies with paired germline and somatic data. If this observation holds in future studies, the lack of inter-ancestry variability encouragingly implies that diagnostic and therapeutic efforts directed towards oncogenic genes may be applicable for all populations.

As with most large-scale genomic studies of pooled patient cohorts, our analyses have several important limitations that present opportunities for future studies. First, we focused on point mutations and small indels, leaving copy number, structural, and epigenomic alterations unexplored. Integrating these data could provide a more complete portrait of tissue-specific oncogenesis, especially for tumor suppressor genes (25). Second, we relied on GTEx bulk RNA-seq profiles to estimate baseline gene expression, which reflect heterogeneous mixtures of cell types and may not capture the precise cell(s) of origin; integrating single-cell or lineage-resolved transcriptomic data could provide more accurate insights (26). Third, despite the large cohort size of AACR Project GENIE, statistical power remained limited for ultra-rare cancers and for analyses stratified by race or ethnicity, particularly among non-White populations. Expanding to ancestrally diverse cohorts will be essential for identifying any existing population-specific drivers of rarer cancers. Fourth, our gene-level analysis did not account for allelic heterogeneity among variants within the same gene—for example, different oncogenic alleles of *KRAS* (*e*.*g*., G12D, G12V) are known to have distinct signaling consequences and tissue-dependent activity (27). Together, these considerations underscore the conservatism of our current findings while highlighting clear paths toward a more mechanistic and clinically actionable understanding of tissue-specific oncogenesis.

In summary, by systematically quantifying tissue-restricted patterns of functional somatic mutations, our study challenges the implicit multi-cancer framing of some driver gene catalogs and emphasizes the value of a subtype-resolved perspective. This approach expands the list of potential drivers across the cancer spectrum and also provides a quantitative scaffold for investigating the biological and clinical consequences of tissue-specific oncogenesis.

## Materials and Methods

### Data curation

We analyzed somatic mutation data from the American Association for Cancer Research (AACR) Project GENIE v18.0 (4), which comprised 250,018 tumor samples from 211,526 patients across 835 cancer subtypes. To avoid ambiguity in tissue-of-origin and to reduce patient duplication due to serial biopsies, we first excluded all metastatic samples before retaining one representative sample per patient (prioritizing primary tumors; see Supplementary Methods).We further collapsed all OncoTree subtypes to level 3 before discarding tissues/subtypes that were underpowered; specifically, we excluded all subtypes that represented <0.1% of the overall GENIE cohort, <1% of all samples within their parent tissue, or had <1,000 samples overall. A site bias–flagging step (see Supplementary Methods) removed 1,900 likely misannotated samples, yielding a final cohort of 146,394 primary tumors spanning 265 subtypes (Dataset S01).

### Definition of subtype-resolved gene mutation frequencies

We first grouped all somatic mutations by predicted functional consequence (*e*.*g*., missense, LoF, silent). For each gene in each cancer pair, raw mutation frequencies were defined for each gene in each cancer type as the fraction of all samples with at least one mutation reported divided by all samples in that cancer type that had any sequencing coverage for the gene. We subsequently normalized these frequencies for cancer-type-specific mutational burden and gene-level mutability, the latter corrected using gnomAD-derived mutation rates adjusted for coding sequence length (28). Statistical enrichment was quantified by chi-squared tests, in which the observed counts for each gene-cancer pair were compared to expected counts derived from a gene length-adjusted background within the same cancer subtype accounting for subtype-specific exome-wide mutational density. We assessed all chi-squared tests at Bonferroni-adjusted significance and further imposed a fold-change cutoff (≥10-fold enriched vs. baseline), ensuring that detected events represented potent, major mutational drivers of each subtype rather than genes with weaker effects situationally driven to significance by variation in sample size across subtypes. Refer to Supplementary Methods for details of secondary analyses based on this dataset, such as epidemiologic incidence adjustment of mutation frequencies or comparison of frequencies between self-reported race/ethnicity groups.

### Computation of tissue specificity scores

To quantify how selectively each gene is mutated across cancer types, we defined a tissue specificity score for each gene by comparing the number of significantly enriched cancer types to the number of adequately powered cancer types for that gene. Statistical power for each gene-cancer pair was estimated using a noncentral chi-squared model parameterized by expected effect sizes from the IntOGen driver gene compendium. Expected effect sizes were assigned as OR=9.7 (combined), OR=16.4 (LoF), and OR=13.2 (missense) relative to baseline. (see Supplementary Methods) (6). In this framework, power reflects whether the number of sequenced samples for a given gene-cancer pair is sufficient to detect enrichment, assuming a plausible driver-like effect size. Cancer types with power >0.5 were considered adequately powered. We then computed a tissue specificity score for each gene as the number of well-powered genes divided by the number of significantly mutated genes after pruning direct parent-child OncoTree relationships to avoid double-counting closely related cancer types. Finally, we scaled these scores from 0 to 100, with higher values reflecting greater tissue specificity.

### Integration of normal tissue expression and interaction networks

To assess whether tissue specificity of mutations related to baseline gene expression, we integrated GTEx v10 median TPM values across 23 tissues mapped to the top-level OncoTree nodes that also were included in our filtered version of the GENIE dataset (14). Expression breadth was defined as the number of tissues in which a gene was expressed above 5 TPM (Fig. S6). We correlated expression breadth with tissue specificity scores using Spearman and linear regression, stratifying missense genes by oncogene/TSG status based on COSMIC annotations (15). We next identified “peripheral” genes for each significantly mutated gene in these analyses. Peripheral genes were defined as those that were differentially expressed between mutationally permissive vs. non-permissive tissues for each cancer gene in question. Differential expression was tested using either t-tests or z-tests across 18,772 protein-coding genes defined by GENCODE v48 that were also quantified in GTEx (see Supplementary Methods) (29). To test whether these peripheral genes were functionally linked to anchor genes, we assessed their enrichment in PPI networks derived from IntAct (21), testing enrichment across 1-4 degrees of network proximity using Fisher’s exact test.

## Acknowledgments

We thank Sasha Gusev and Eli Van Allen for their input and guidance. We would also like to acknowledge the American Association for Cancer Research and its financial and material support in the development of the AACR Project GENIE registry, as well as members of the consortium for their commitment to data sharing. Interpretations are the responsibility of study authors. This work was supported by an award from the Cancer Research UK Grand Challenge and the Mark Foundation to the SPECIFICANCER team and NCI K99 CA286805 to RLC. KMH is also supported by the Dana-Farber Cancer Institute Hale Center for Pancreatic Cancer Research and the Project P fund. RLC received additional support from the Rossy Foundation Fund at Myriad Canada (formerly KBF Canada).

