## Supporting Information for "The tissue specificity of cancer genes"

**This PDF file includes:**

Materials and Methods
Figures S1 to S8
Legends for Datasets S1-S3
SI References

**Other supporting materials for this manuscript include the following:**

Datasets S1-S4

### Supporting Information Text
### Materials and Methods

#### Data acquisition and processing

Data was acquired from the American Association for Cancer Research (AACR) Project GENIE, version 18.0-public release, comprising 250,018 tumor samples from 211,526 patients across 835 cancer subtypes annotated using OncoTree ontology (1, 2). Clinical sample metadata were retrieved from *data_clinical_sample.txt*.

Samples labeled as "metastatic" (n = 70,639; 28.3%) were excluded to avoid ambiguity in tissue-of-origin. To prevent overrepresentation of individual patients, samples were deduplicated by selecting a single representative sample for each unique patient ID using the following criteria: preference for primary tumors, exclusion of samples with unknown tumor type, oldest sample by age at diagnosis, and random selection in case of ties. This filtering procedure yielded a cohort of 151,915 unique patient samples covering 820 distinct cancer subtypes.

To reduce noise from sparsely sampled groups, samples were collapsed to OncoTree tissue-level categories. OncoTree nodes representing less than 0.1% of the overall GENIE cohort (≤151 samples) were excluded. To further control for subtype granularity, OncoTree subtypes were collapsed up to level 3 terms and subsequently retained only if they represented ≥1% of patients within their parent tissue or ≥1,000 patients overall. This dual criterion ensured removal of underpowered subtypes while retaining clinically relevant & well-powered subtypes within high-incidence cancers (e.g., lung, bowel). Samples labeled "Unknown" were excluded, yielding a cohort of 148,294 unique patient-samples.

#### Somatic mutation annotation and filtering

Somatic mutation calls were obtained from *data_mutations_extended.txt* and grouped into three functional classes based on *Variant_Classification*: missense (Missense_Mutation, In_Frame_Ins/Del), LoF (Nonsense_Mutation, Splice_Site, Frame_Shift_Del, Frame_Shift_Ins, Nonstop_Mutation, Splice_Region, Translation_Start_Site), and silent ('Flank, 3'UTR, 5'Flank, 5'UTR, Intron, RNA, Silent). Deduplicating mutation calls after regrouping yielded 797,650 missense, 284,969 LoF, and 200,802 silent mutations.

To correct for site-specific biases in cancer type annotation, we applied a center-level quality control filter. For each (gene, cancer, center) tuple, we compared the mutation frequency to the background frequency pooled across all other centers using Fisher's exact test, performed separately for LoF and missense mutations. Tuples with $P < 0.05$ and mutation frequency fold-change (FC) >5 compared to all other centers were flagged and removed. An additional FC filter on center-specific mutation frequencies minimized false positives from high-powered centers and reduced correlation between flagged events and center size. A total of 69 biased (gene, cancer, center) tuples were removed. In total, this procedure led to the exclusion of 1,900 likely misannotated samples, reducing the overall cohort from 148,294 to 146,394 samples spanning 265 distinct cancer subtypes (Dataset S01, patient count per cancer subtype). After filtering, the final curated GENIE dataset contained 752,971 missense, 263,230 LoF, and 139,755 silent mutations, which were subsequently tallied for each gene and cancer type separately for missense alone, LoF alone, and all nonsynonymous mutations (LoF + missense).

#### Sequencing coverage matrix construction

To account for gene- and cancer-type–specific sequencing depth, we constructed a denominator matrix recording how often each gene was sequenced in each cancer type. Assay IDs from *data_clinical_sample.txt* were matched to gene content in *genomic_information.txt* to determine panel coverage per sample. The resulting matrix (1,579 genes × 265 cancer types) aligned with the dimensions of the mutation matrices.

### Mutation matrix filtering for downstream analysis

We next wanted to reduce noise in our downstream analyses contributed by undersequenced—and therefore underpowered—genes. To achieve this, we first computed the binomial 90% confidence interval (CI) for mutation frequencies for each gene–cancer pair using the *binconf()* function from the *Hmisc* R package. Genes were retained if the maximum 90% CI lower bound of mutation frequencies across any cancer type was ≥10% in at least one of the three mutation matrices (LoF, missense, or combined). This step ensured the inclusion of genes that were both sufficiently mutated and sequenced to support reliable mutation frequency estimates. A total of 590 genes across 265 cancer types passed this filter and were used in downstream analyses (Dataset S02).

### TCGA mutation frequencies

For comparison, we also curated somatic mutation data from TCGA as follows: mutations were grouped into missense, LoF, and silent categories following the same classification scheme as for GENIE (3). After deduplication and restricting to solid tumors, 127,506 missense or LoF mutations from 9,572 samples were retained. Cancer types were assigned using clinical metadata *clinical_PANCAN_patient_with_followup.tsv*. For pan-cancer mutation frequencies, each gene's mutation count was defined as the number of samples harboring at least one missense or LoF mutation, using total number of patients (n=9,722) as the denominator. All genes were assumed to be sequenced in all samples given that TCGA applied whole-exome sequencing as opposed to targeted gene panels (Dataset S03).

### SEER incidence-adjusted mutation frequencies

To approximate population-level mutation burden, tissue-level mutation frequencies from GENIE and TCGA were weighted by SEER-reported incidence proportions (Dataset S03) (4). We restricted this analysis to solid tumors due to the technical challenges in detecting low-frequency clones in liquid tumors and the greater heterogeneity of normal and cancer cells in liquid biopsies. Tissue labels were harmonized using OncoTree annotations, with 27 SEER sites mapped to 19 GENIE tissue-level solid cancer types. Separately, 91 TCGA tissue-cancer type combinations were mapped to 23 GENIE tissue types, 17 of which also matched to SEER. After restricting to these 17 shared solid tissues, 8,889 TCGA samples with 124,782 mutations were used to compute tissue-specific mutation frequencies for adjustment. Plotting was restricted to genes sequenced ≥100 times across all cancer types and present in ≥10% of cancer types (n = 552 of 590), with a requirement of ≥30 sequenced samples per gene-cancer pair and was limited to driver genes annotated in IntOGen (n = 225). We observed that some genes had higher SEER-adjusted mutation frequencies compared to their GENIE or TCGA unadjusted frequencies. This pattern reflects differences in calculation: SEER-adjusted frequencies are obtained by summing tissue-specific mutation frequencies weighted by cancer incidence, whereas pan-cancer frequencies are calculated from aggregated mutation counts across all tumors without incidence weighting. Consequently, genes with high mutation frequencies in a small number of high-incidence cancers can appear artificially elevated after adjustment. To prevent misclassification of such cases as pan-cancer outliers, we excluded these genes from downstream interpretation.

### Normalization for mutation burden and gene length bias

We applied a chi-squared based normalization procedure to each gene in each cancer type to account for variation in subtype-specific genome-wide mutation burden and gene-specific differences in mutability. For each cancer type, the observed mutation counts per gene were compared to a background model derived from the mutation frequencies of all other sequenced genes in that cancer type. Gene length is a major confounder in statistical enrichment testing of gene-based mutation burden. Further, it is well established that there is additional variability in gene mutation rates even among genes of the same length, which is at least partially attributable to the differences in mutability of the primary nucleotide composition within the coding sequences (CDS) of each gene (5). Gene CDS lengths were computed from *genomic_information.txt*, which records the start and end positions and exon status of each mutated gene included in each gene panel. For each gene, a single length value was calculated by averaging across all panels in which the gene was covered. Among 9,321 mutation calls from Wake Forest Baptist Medical Center, 9,312 had identical start-end positions, while 47 lacked coverage in both exonic and intergenic regions; these samples were excluded from mean CDS calculations. Given that these gene-specific mutation rate estimates were positively correlated with gene CDS length (Fig. S7), we simultaneously corrected for biases in gene length and mutation rate using a scalar derived from gnomAD mutation frequencies, calculated as the ratio of a gene's gnomAD-based mutation frequency to the background mean frequency across all genes sequenced in each cancer type (6). For genes lacking gnomAD data ("SSX2" and "HMGN2P46" with no LoF or missense records; "RPA4" and "OR5L1" with no LoF records), we used a linear model of mutation frequency versus GENIE-reported CDS length to impute values. We implemented these gene length and mutation rate adjustments in our chi-squared tests to determine mutational significance by multiplying each gene's background mutation frequency by the corresponding scalar, ensuring that expected mutation counts were adjusted for intrinsic length-related mutability.

We also recorded FC per gene-cancer pair, defined as the observed mutation frequency of the gene divided by the background mutation frequency in that cancer type. This FC quantifies effect size and complements statistical significance in downstream filtering. The procedure yielded three chi-squared matrices and three FC matrices (missense, LoF, and combined), reflecting mutation enrichment relative to a gene length-adjusted background.

**Binary mutation enrichment matrices**

To classify gene-cancer pairs as significantly enriched or not, we binarized the chi-squared results using a Bonferroni-corrected significance threshold. Specifically, we computed the critical chi-squared statistic corresponding to a Bonferroni-adjusted *p*-value cutoff of 0.05 ($P < 1.60 \times 10^{-7}$) and marked pairs exceeding this threshold as statistically significant.

An important *post hoc* step in our determination of mutational significance was to impose a uniform minimum FC across all genes and cancer types, which was designed to reduce bias from cancer type prevalence—where common cancers have relatively greater power to detect comparatively more modest enrichments. We used an Empirical Bayes approach for defining a minimum FC that corresponded to existing sets of established, biologically meaningful cancer driver genes. Specifically, we examined the distribution of FC values across all significantly enriched (gene, cancer) and selected the cutoff at the inflection point of this distribution (Fig. S8). We integrated this minimum FC alongside our P-value criteria as follows: a gene was considered significantly enriched in a cancer type only if it exhibited (1) a chi-squared statistic exceeding the Bonferroni-derived critical value as above and (2) a FC >10. This dual threshold ensures that identified enrichments are both statistically robust and of sufficient effect size, minimizing the detection of weak effects in highly powered cancers while preserving strong signals in rarer subtypes. Indeed, we observed that this post hoc FC filter dramatically improved comparability in the counts of significant drivers between cancer types, especially between rare and common cancers (Fig. S4).

**Gene mutation pattern stratified by self-reported race/ethnicity**

To assess mutational enrichment across racial and ethnic groups, we repeated the binary mutation enrichment analysis described above, this time stratified by self-reported race and ethnicity in GENIE. Samples labeled as "Spanish/Hispanic" ethnicity were classified as "Spanish/Hispanic," (n=9,472) while all remaining samples were grouped using the race field into "Asian" (n=6,502), "Black" (n=6,365), or "White" (n=76,509). We emphasize that these categories reflect the available self-reported information recorded in GENIE and were not modified or inferred further; we do not interpret these to be perfect proxies of underlying genetic ancestry but enabled an initial coarse grouping of patients based on their self-reported demographics.

For each self-reported demographic group defined above, we constructed mutation count matrices, computed chi-squared statistics and fold changes, and applied significance thresholds based on a Bonferroni-adjusted *p*-value cutoff of 0.05, using the number of tests specific to each (ethnicity, mutation consequence) combination. This corresponded to $p < 7.01\text{x}10^{-6}$ for Asian, $8.60\text{x}10^{-6}$ for Black, $3.52\text{x}10^{-6}$ for Hispanic, and $1.32\text{x}10^{-6}$ for White. An FC > 10 threshold was then applied, yielding binary matrices for missense, LoF, and combined mutations. Gene-cancer-consequence combinations were evaluated by binomial test if the "White" binary matrix entry was 0 while the corresponding entry in any other group was 1. To assess group-specific enrichments, we performed binomial tests comparing each non-White group to the White population baseline.

**Identifying cancer types with adequate power for each gene**

For each gene-cancer pair, we assessed whether there was sufficient statistical power to detect mutation enrichment. We modeled the alternative hypothesis as a noncentral chi-squared distribution with 1 degree of freedom and a noncentrality parameter derived from an expected effect size.

To parameterize the expected effect size, we used odds ratios (ORs) derived from the IntOGen cancer driver gene compendium (filtered at combined Q-value < 0.05 and >1% of samples affected in the cohort) (7). For a given gene with total sample size N, expected counts for mutated and non-mutated samples (adjusted for gene length using the pre-computed correction scalar) were denoted $E_{mut}$ and $E_{wt}$, respectively. Given an expected OR, we computed the target ratio of observed mutated to wild-type samples as:

$$R = \frac{O_{mut}}{O_{wt}} = OR \times \frac{E_{mut}}{E_{wt}}$$

Solving the system $O_{mut} + O_{wt} = N$, we obtain:

$$O_{mut} = \frac{R \times N}{1 + R}, O_{wt} = N - O_{mut}$$

These values were used to calculate the noncentrality parameter (NC) of the chi-squared distribution:

$$NC = \frac{(O_{wt} - E_{wt})^2}{E_{wt}} + \frac{(O_{mut} - E_{mut})^2}{E_{mut}}$$

The *pchisq()* function in R was used to calculate the power as the probability of observing a chi-squared statistic greater than the critical value under the null, given the assumed noncentral alternative. Power was then computed as:

$$Power = 1 - P(\chi_1^2 < X_c \mid NC) = 1 - pchisq(X_c, df = 1, ncp = NC)$$

In plain terms, this approach asked: *Given the number of times a gene was sequenced in this cancer type, do we have enough power to detect enrichment assuming a biologically meaningful effect size?* We defined a cancer type as "powered" for a given gene if this *post hoc* power exceeded 0.5. This step ensured that tissue specificity scores are not inflated by situations where a gene was simply not sequenced enough in some cancer types to provide a confident mutation frequency estimate.

**Specificity score definition**

We computed tissue specificity scores for each gene as a fraction comparing significantly mutated cancer types to all powered cancer types. The numerator of the calculation for each gene was taken as the number of "powered" cancer types defined above. The denominator for each gene was assigned as the number of cancer types in which each gene was significantly enriched for mutations. Formally, we expressed these scores as the following ratio:

$$Specificity\ Score_{raw} = \frac{\#Powered\ Cancer\ Types}{\#Significantly\ Mutated\ Cancer\ Types}$$

This ratio captured how uniquely a gene was mutated after accounting for the high degree of gene and cancer type heterogeneity in sequencing coverage and sample sizes. A high score indicates that a gene is mutated in a small subset of cancer types despite sufficient power to detect enrichment in many, suggesting true tissue specificity.

To avoid score deflation due to OncoTree's hierarchical ontology, we excluded parent cancer types when a gene was significantly mutated in both parent and immediate child. This was applied to both the powered (numerator) and enriched (denominator) cancer type sets, under the assumption that the parent signal was not independent from the child and offered less granularity.

After adjusting these scores for parent-child redundancy, we scaled the final tissue specificity scores from 0 to 100 for ease of interpretability, with higher scores indicating greater subtype specificity. Percentile ranks were also computed for comparison (Dataset S04). Not all genes were significantly enriched in any cancer subtypes or did not have adequate power in any subtypes to meet our criteria above; due to these limitations, we were able to compute missense specificity scores for 56 genes, LoF scores for 54 genes, and 73 for either class.

**Expression data integration**

We used GTEx Analysis V10 median gene-level TPM values (RNASeQC v2.4.2) to assess gene expression breadth (8). Of 68 GTEx tissues, 23 were mapped to 31 GENIE tissue-level OncoTree categories. For tissues with duplicate mappings, TPMs were converted to fractions (divided by 1,000,000) and harmonically averaged.

We binarized these expression quantifications per tissue where genes with ≥5 TPM were classified as meaningfully expressed, and all genes with <5 TPM were labeled as not expressed. Expression breadth was defined as the number of GTEx tissues (out of 23) in which a gene was expressed. While we acknowledge that this ≥5 TPM threshold is arbitrary, we note that this threshold is commonly used in other transcriptomic analyses, and we found that the exact selection of TPM threshold did not meaningfully impact our downstream analyses (Fig. S6)

We assess the relationship between expression breadth and normalized tissue specificity scores for each mutation class (LoF, missense, combined) with Spearman correlation tests. Linear regressions were also performed. Missense genes were further stratified into oncogenes and TSGs using COSMIC annotations (9).

**Defining genes peripheral to each significantly mutated cancer gene**

Peripheral genes were defined as those with tissue-specific expression differences in normal tissues that tracked with the somatic mutation profile of any given cancer ("anchor") gene. For each anchor gene, tissues were divided into two groups: (1) *permissive* tissues, in which the anchor gene was significantly enriched for functional mutations, and (2) *non-permissive* tissues, in which no such enrichment was detected. Expression levels of 18,772 protein-coding genes—defined by the intersection of GTEx expression data with GENCODE v48 basic annotations—were compared between the two tissue groups (10). Two-sided *t*-tests were applied when both groups contained more than one tissue; otherwise, a *z*-test was used. Genes with FDR-adjusted $p < 0.05$ were classified as *peripheral genes*, indicating tissue-specific differential expression in relation to the anchor gene's mutation profile.

**Defining a human protein-protein interaction network**

We built a network of human protein-protein interactions (PPIs) based on data available from the Ensembl IntAct database (11). We downloaded all molecular interactions asserted in IntAct as of December 5, 2024, involving two or more human proteins with gene symbols or aliases that matched canonical protein-coding genes also defined in the Gencode human gene reference (v47) (10). To ensure we only retained high-confidence direct interactions, we excluded all interactions determined by "physical association," "colocalization," "proximity," or "association"; upon review, all remaining terms indicated likely direct PPIs. We next constructed a PPI network from this curated dataset with proteins were nodes and interactions as edges. Lastly, we pruned any edges from this network present in the IntAct "negative" interaction files, which contain putative interactions that have been convincingly refuted by published evidence after curation by the IntAct team. All remaining edges in this graph were considered PPIs for our downstream analyses in this study.

**Peripheral gene enrichment in anchor gene PPI networks**

We intersected the sets of peripheral genes defined from normal tissues from GTEx with the anchor gene-centric PPI networks curated from IntAct, as described above. We tested for enrichment of differentially expressed peripheral genes within each PPI network degree (1–4) by intersecting proteins within that network radius with the anchor's peripheral gene set. We subsequently performed Fisher's exact tests to determine enrichment of peripheral genes among the interaction partners. ORs and *p*-values were calculated for each degree.

**Figures**

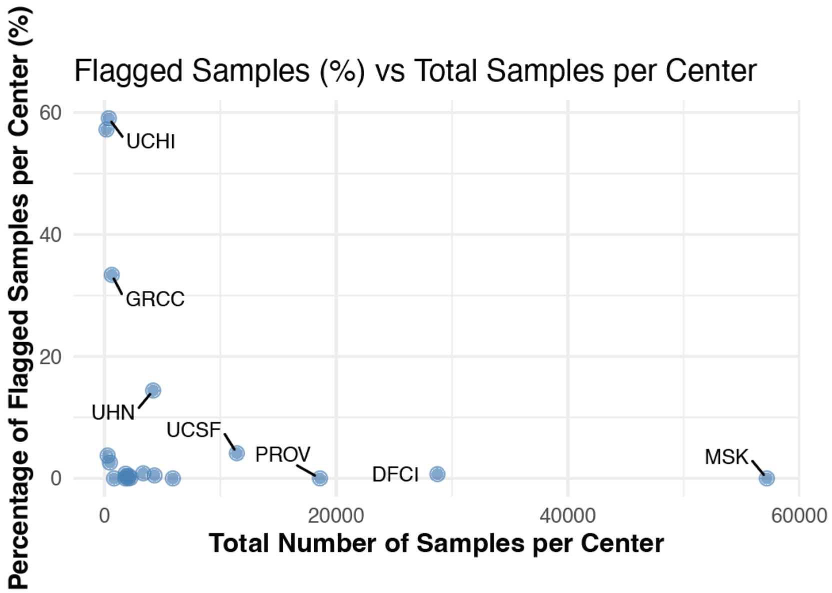


**Fig. S1.** Flagged vs. total samples per sequencing center.

We found that certain centers had higher rates of likely sample misannotation, which were flagged and removed from our analyses. Here, we compare the total number of samples submitted (x-axis) and the number of samples flagged as likely misannotated (y-axis). Each point represents one center, with labels indicating center names.

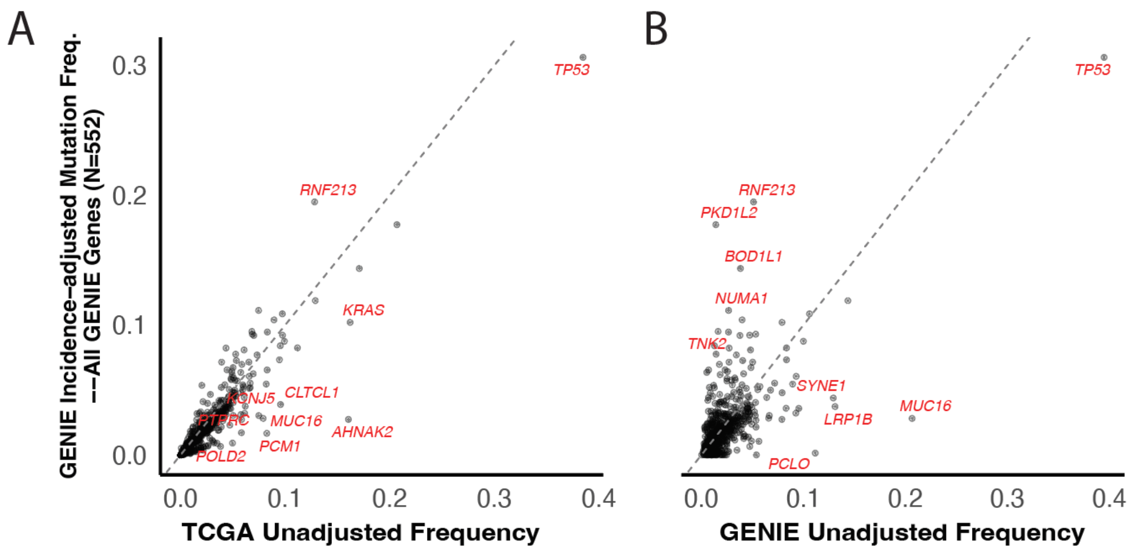


**Fig. S2.** Comparison of incidence-adjusted mutation frequencies with unadjusted and TCGA-based estimates.

Scatterplots show gene mutation frequency with SEER adjustment for GENIE versus mutation frequency without SEER incidence adjustment for (A) GENIE and (B) TCGA. Dashed lines represent linear fits.

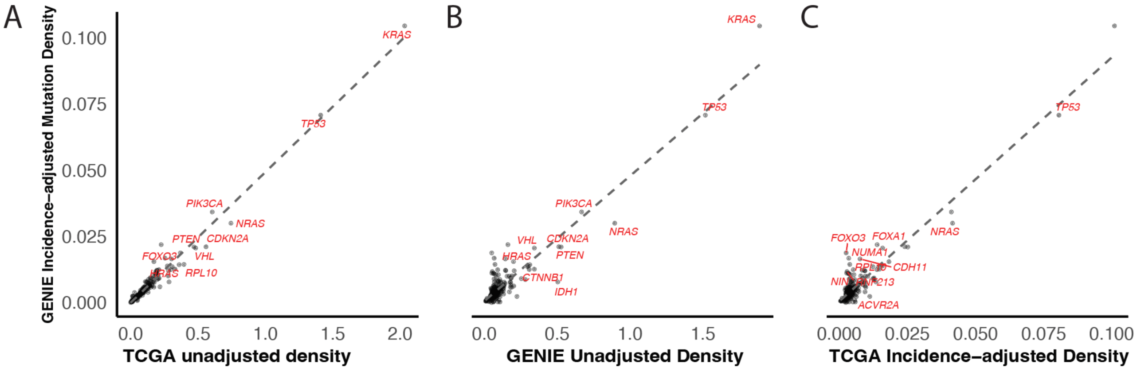


**Fig. S3.** Comparison of incidence-adjusted mutation densities with unadjusted GENIE and TCGA, and adjusted TCGA estimates.

Gene length-normalized mutation densities from GENIE (y-axis, SEER incidence-adjusted) compared with three reference estimates (x-axis): unadjusted GENIE frequencies, unadjusted TCGA frequencies, and TCGA SEER-adjusted densities. Dashed lines represent linear fits. Genes with the largest discrepancies between incidence-adjusted and reference values are labeled with red text.

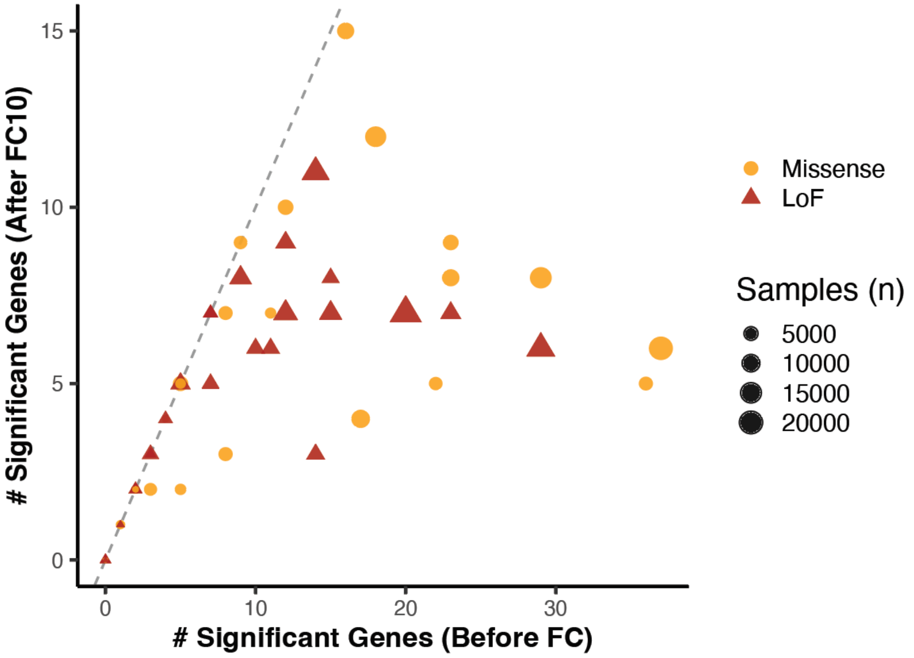


**Fig. S4.** Comparison of significant driver gene counts before and after fold-change correction.

We compared the number of significant missense (orange circles) and LoF (red triangles) genes identified per tissue before (x-axis) and after (y-axis) applying the ≥10-fold enrichment correction. Point size is proportional to the number of samples sequenced in each tissue. The dashed diagonal line is a visual guide for $y = x$. Most points fall below the diagonal, reflecting a reduction in the number of significant genes after correction, particularly in well-sampled tissues.

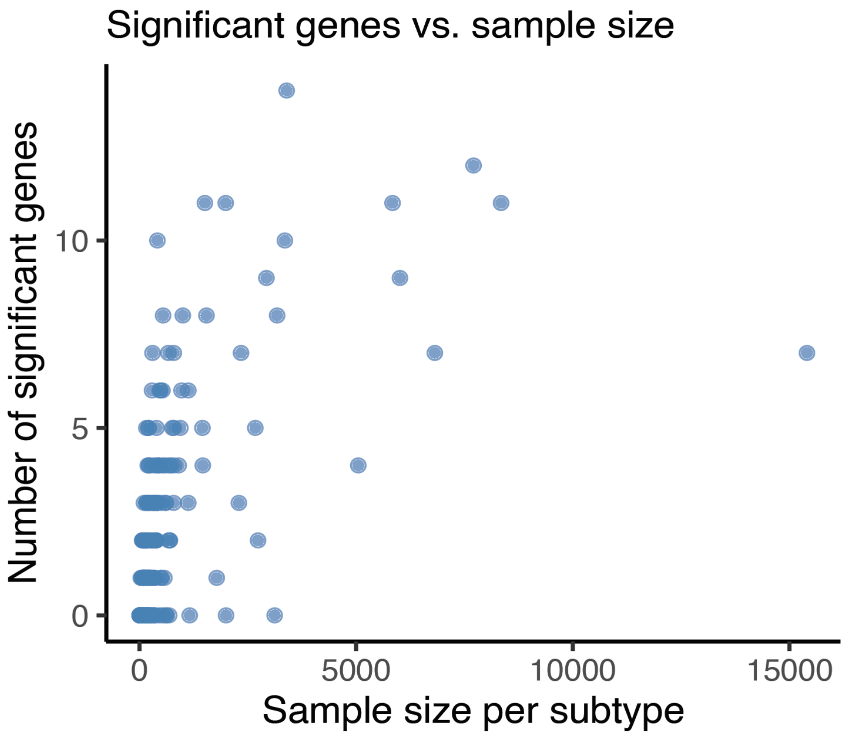


**Fig. S5.** Relationship between cancer sample size and number of significant genes per cancer subtype after applying ≥10-fold enrichment correction.

Scatterplot showing the number of significantly mutated genes (y-axis) vs. sample size (x-axis) across all cancer subtypes. Each point represents a single OncoTree subtype. The slight positive association highlights the influence of sample size on power to detect significant gene-subtype enrichments, even after fold-change-based filtering.

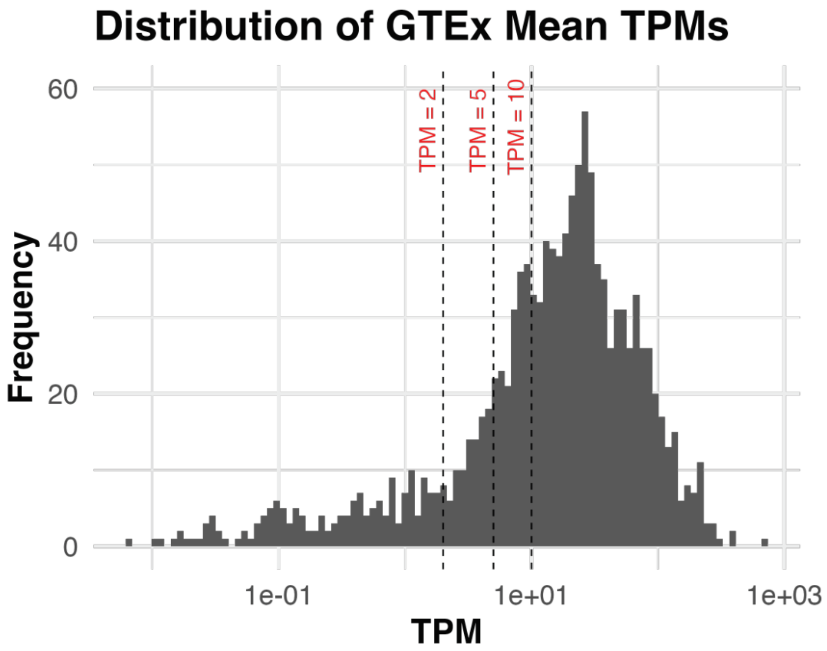


**Fig. S6.** TPM expression distribution for the 590 genes considered in our final analyses.

Varying the threshold across a reasonable range (2–10 TPM) yielded highly similar distributions of expression breadth.

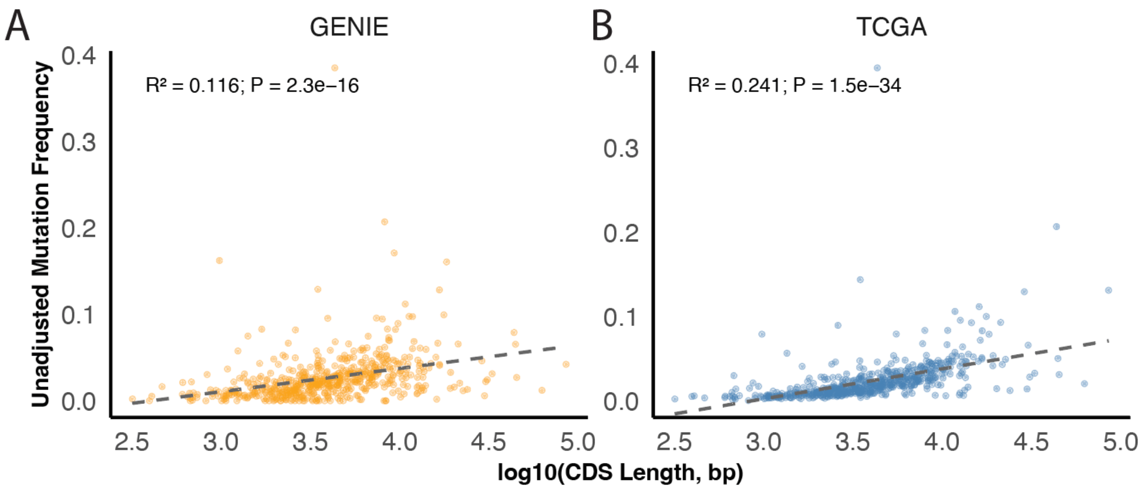


**Fig. S7.** Comparison of gene length vs. unadjusted pan-cancer mutation frequencies.

As expected, we found that gene length was strongly correlated with mutation frequencies in both the GENIE and TCGA datasets. Here, we provide $\log_{10}$(CDS length, bp) versus gene mutation frequency without SEER incidence adjustment for (A) GENIE and (B) TCGA for all genes in our analyses. Dashed lines represent linear fits.

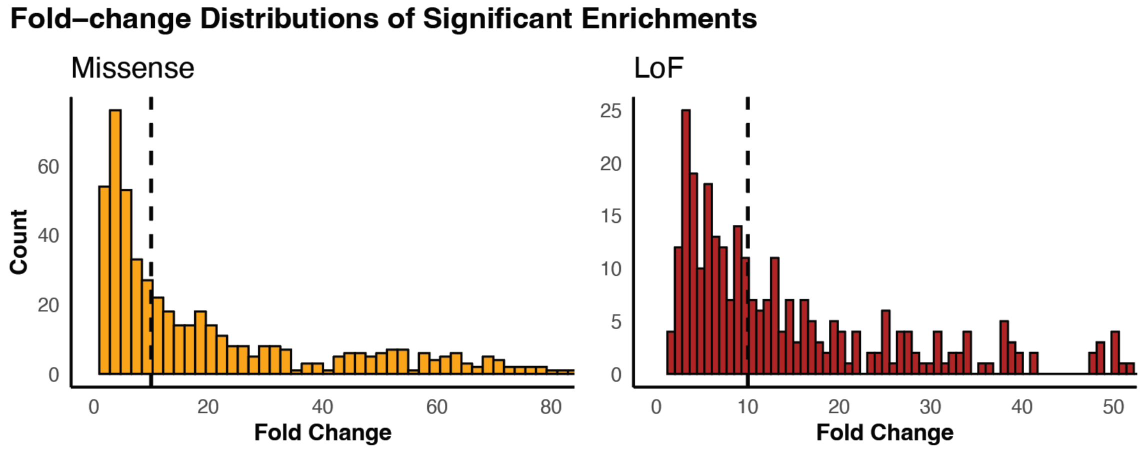


**Fig. S8.** Distribution of FC values for mutationally enriched gene-cancer pairs, shown separately for missense mutations (orange) and LoF mutations (red). Axes are bounded to ranges that capture the inflection point of each distribution (0–80 for missense, 0–50 for LoF). The dashed line marks FC = 10, the inflection point of the distribution, which was the chosen threshold for downstream analyses.

**Dataset S1 (separate file).** Sample count by cancer subtype. Number of primary tumor samples for each of the 265 cancer subtypes included in the final analysis. Subtypes are annotated using OncoTree labels.

**Dataset S2 (separate file).** Mutation frequency matrices (590 genes x 265 cancer subtypes) for (1) combined LoF or missense, (2) LoF, and (3) missense mutations.

**Dataset S3 (separate file).** A set of 553 genes from the curated list of 590, restricted to those sequenced at least 100 times across all cancer types. For each gene, we provide the CDS length (averaged across all gene panels that include the gene), along with raw mutation frequencies, SEER-incidence-projected mutation frequencies, and length-adjusted mutation densities (with and without SEER projection) from both GENIE and TCGA.

**Dataset S4 (separate file).** Cancer gene lists ordered by tissue specificity score.
Reported separately for (1) combined LoF or missense, (2) LoF, and (3) missense mutations. Within each list, genes are ordered from highest to lowest tissue specificity score, with higher scores indicating stronger subtype restriction.
